# The impact of genetic background and sex on the phenotype of IL-23 induced murine arthritis

**DOI:** 10.1101/2021.02.03.429523

**Authors:** Emma Haley, Mederbek Matmusaev, Imtiyaz N. Hossain, Sean Davin, Tammy M. Martin, Joerg Ermann

## Abstract

**Background:** Overexpression of IL-23 in adult mice by means of hydrodynamic tail vein injection of IL-23 minicircles has been reported to result in spondyloarthritis-like disease. The impact of genetic background and sex on the disease phenotype in this model has not been investigated.

**Methods:** We compared male B10.RIII mice with male C57BL/6 mice, and male with female B10.RIII mice after hydrodynamic injection of IL-23 enhanced episomal vector (EEV) at 8-12 weeks of age. We monitored clinical arthritis scores, paw swelling, and body weight. Animals were euthanized after two weeks and tissues were harvested for histology, flow cytometry and gene expression analysis. Serum cytokine levels were determined by ELISA.

**Findings:** Male B10.RIII mice developed arthritis in the forepaws and feet within 6 days after IL-23 EEV injection; they also exhibited psoriasis-like skin disease, colitis, weight loss, and osteopenia. In contrast to previous reports, we did not observe spondylitis or uveitis. Male C57BL/6 mice injected with IL-23 EEV had serum IL-23 levels comparable with B10.RIII mice and developed skin inflammation, colitis, weight loss, and osteopenia but failed to develop arthritis. Female B10.RIII mice had more severe arthritis than male B10.RIII mice but did not lose weight.

**Conclusions:** Systemic IL-23 overexpression results in spondyloarthritis-like disease in B10.RIII mice. The development of extra-articular manifestations but absence of arthritis in C57BL/6 mice suggests organ-specific genetic control mechanisms of IL-23 driven inflammation. Discrepancies regarding the phenotype of IL-23 induced disease in different labs and the sexual dimorphism observed in this study warrant further exploration.

## Introduction

Spondyloarthritis (SpA) is a family of inflammatory rheumatic diseases including ankylosing spondylitis (AS) and psoriatic arthritis (PsA). 0.5-1.5% of the adult population in the United States have SpA [1]. Enthesitis, inflammation at the attachment sites of tendons and ligaments to bone, is thought to be the key process driving inflammation in the spine and peripheral joints in SpA [2]. Inflammation and tissue damage may also occur outside of the musculoskeletal system. Commonly affected organs are the skin (psoriasis), eye (uveitis) and gut (inflammatory bowel disease) [3]. Except for AS, which is more common in males (male:female ratio 2-3:1), SpA is equally common in men and women. However, extra-articular manifestations, such as psoriasis, colitis, and enthesitis occur at a higher frequency in females with AS, and females with AS or PsA tend to have higher disease activity [4, 5]. The mechanisms behind the sexual dimorphism in SpA are incompletely understood.

Multiple lines of evidence including genome-wide association studies, animal models, and therapeutic clinical trials, support a critical role for the IL-23/IL-17A pathway in the pathogenesis of SpA [6, 7]. For instance, polymorphisms in multiple genes involved in IL-23 receptor signaling such as *IL12B, IL23R, JAK2* and *TYK2* have been shown to be associated with AS, and polymorphisms in the *IL23R* gene are associated with PsA [8, 9]. IL-23 is produced by myeloid cells, such as dendritic cells and macrophages, and acts on Th17 cells and a variety of innate lymphocytes to produce IL-17A, IL-17F, IL-22 and GM-CSF, all of which have been implicated in the pathogenesis of SpA [10–12].

To study disease mechanisms downstream of IL-23, Sherlock et al. hydrodynamically injected mice with IL-23 minicircles. The resulting SpA-like disease was partially mediated by IL-17A and IL-22 and appeared to involve an enthesis-resident IL-23 receptor expressing CD3^+^CD4^-^CD8^-^ T cell population, which produced IL-17A and IL-22 when stimulated with IL-23 *in vitro* [13]. Other investigators showed subsequently that these enthesis-resident double-negative T cells were predominantly γδ T cells [14, 15]. Preliminary experiments in our lab confirmed that B10.RIII mice hydrodynamically injected with IL-23 enhanced episomal vector (EEV), which very efficiently drives hepatic IL-23 production, develop arthritis. However, we noticed phenotypic differences compared with previous publications. Moreover, we found that C57BL/6 (B6) mice, the most widely used inbred strain and genetic background for many transgenic and gene knockout models including IL23R-GFP reporter mice [16], were protected from arthritis development.

The goals of this study were to describe the disease phenotype in susceptible B10.RIII mice injected with IL-23 EEV in our laboratory, compare the disease phenotype in male B10.RIII and B6 mice injected with IL-23 EEV and, in line with the National Institutes of Health (NIH) requirement to consider sex as a biological variable [17], compare the disease phenotype in male and female B10.RIII animals.

## Materials and methods

### Mice

B10.RIII-*H2^r^ H2-T18^b^*/(71NS)*SnJ* mice (abbreviated B10.RIII, Jax #000457) and C57BL/6J (abbreviated B6, Jax #000664) mice were originally obtained from the Jackson Laboratory and maintained in a specific pathogen-free animal facility at Brigham and Women’s Hospital. For all experiments, 8-12 week-old male (body weight 21.6-33.0 g) or female (body weight 17.9-26.0 g) mice were used. Mice were housed under standard 12-hour light/12-hour dark conditions in individually ventilated cages, up to 5 mice per cage, with water and chow (5058 PicoLab Rodent Diet 20) available ad libitum. Littermates were used whenever possible and randomly assigned to experimental and control groups. Data were pooled from multiple independent experiments. All experimental procedures involving animals were approved by the IACUC of Brigham and Women’s Hospital and performed in accordance with federal and institutional guidelines and regulations.

### IL-23 EEV injection and disease monitoring

IL-23 EEV(EEV651A-1, System Biosciences) was hydrodynamically injected in a volume of sterile HBSS (Invitrogen) corresponding to 10% of the animal’s body weight [13]. Mice with failed hydrodynamic injection (<75% of target volume injected) were excluded from the experiment. In most experiments, we administered 500 ng IL-23 EEV per mouse based on preliminary experiments. In a separate dose titration experiment, mice received 500 ng, 50 ng, or 5 ng IL-23 EEV. We monitored the mice for the development of disease every other day, typically in the morning. Each paw was graded for clinical arthritis as follows: 0, normal; 1, minor swelling/erythema; 2, moderate swelling/erythema; or 3, severe swelling/erythema involving the entire paw. Scores from the four paws were summed for a total score from 0 to 12. Paw thickness was measured using calipers. Paw swelling was calculated as the thickness of each paw minus its thickness at baseline, summed for the four paws of each mouse. Disease monitoring was performed by a single observer blinded for group. Animals were euthanized after two weeks by carbon dioxide asphyxiation followed by cervical dislocation. Serum was prepared from heart blood. Spleen and colon weight were recorded. Tissue specimens for gene expression analysis were harvested into RNAlater (Thermo Fisher Scientific).

### Histology

Tissue specimens for histological analysis were fixed in 10% neutral buffered formalin (Sigma) at 4°C for 24 hours. Joint and spine samples were decalcified in 96% formic acid (Electron Microscopy Sciences). Paraffin-embedded sections were stained with hematoxylin and eosin (H&E). Images were captured using a Leica DFC450 digital microscope camera and analyzed with LAS X software (Leica Microsystems).

### Splenocyte Isolation

Single cell suspensions from the spleen were prepared by filtering the disaggregated organ through a 70 μm cell strainer. Cells were further purified with red blood cell lysis buffer (0.15 M NH_4_Cl, 1 mM KHCO_3_, both from Sigma) to eliminate red blood cells. An automated cell counter (Countess, Invitrogen) was used with trypan blue staining to enumerate viable cells.

### Flow cytometry

Surface marker staining was performed in 96-well plates, 2×10^6^ cells per well. First, the cells were stained with fixable viability dye (FVD-450 UV, Thermo Fisher Scientific) in HBSS for 15 minutes. Then the cells were incubated with Fc block (Biolegend) for 10 minutes at room temperature to prevent non-specific antibody binding to Fc receptors. Antibodies for surface markers (Table 1) were added in staining buffer (HBSS with 2 mM EDTA (Boston Bioproducts) and 0.5 % bovine serum albumin (Sigma)) for 30 minutes at 4°C. After washing with staining buffer, the cells were fixed with IC Fixation Buffer (Thermo Fisher Scientific) for 30 minutes at room temperature. An LSRFortessa flow cytometer (BD Biosciences) was used for data collection and FlowJo Software was used for analysis.

**Table 1:**
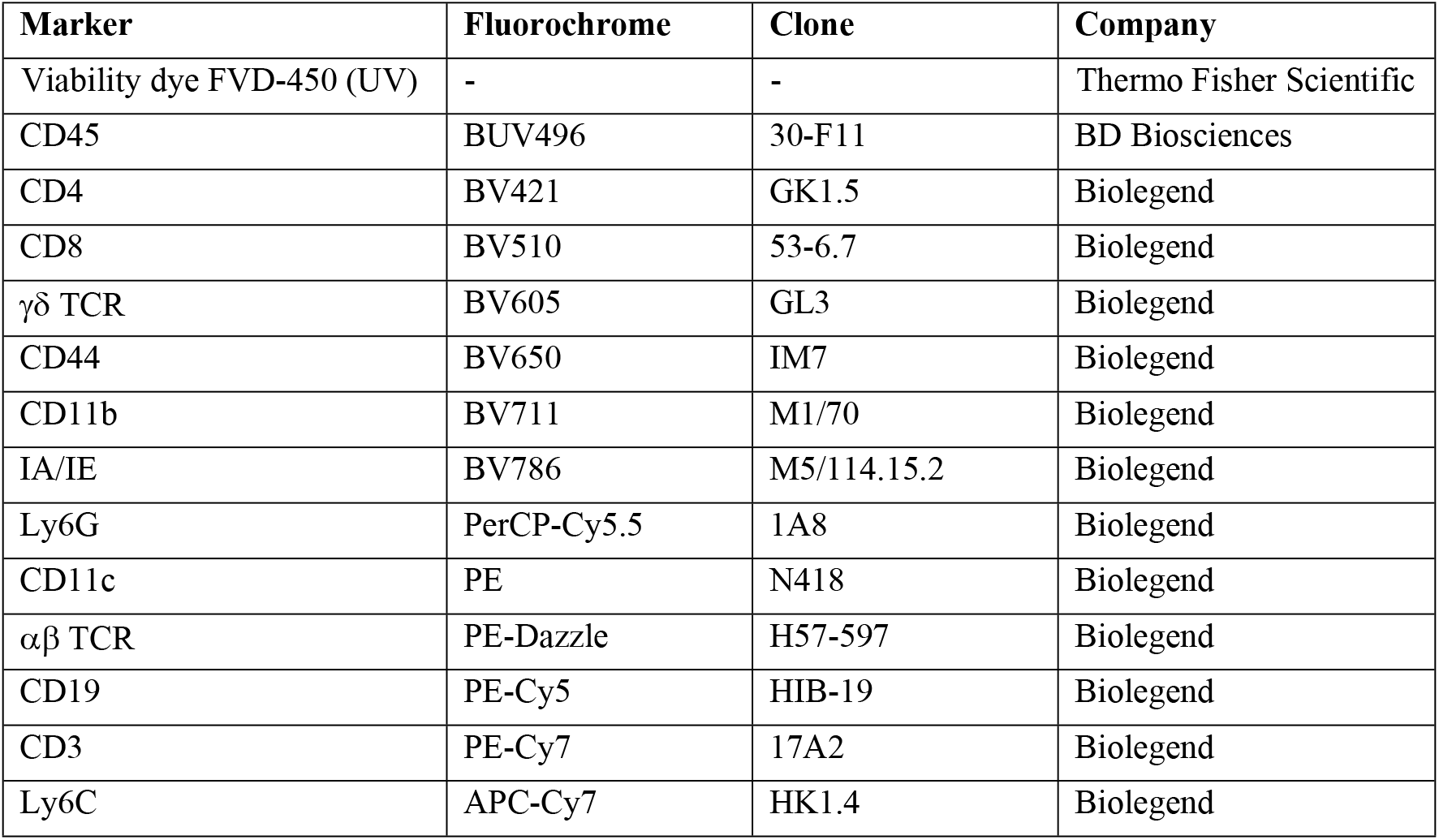
Surface marker antibodies for flow cytometry.

### RNA preparation

Total RNA was prepared using RNeasy mini columns (Qiagen). For RNA preparation from joints and skin, the manufacturer’s protocol was modified as follows: First, the tissue was added to Buffer RLT in Navy Rino Bead Lysis tubes (Next Advance) and homogenized in a bullet blender (Next Advance) for 5-10 minutes at 4°C. The tissue homogenate was cleared by centrifugation, mixed with RNase-free water and proteinase K (100 μg/mL, Sigma) and incubated for 10 minutes at 55°C. All subsequent steps were performed according to the RNeasy mini protocol.

### cDNA Reverse Transcription

The High-Capacity cDNA Reverse Transcription Kit (Thermo Fisher Scientific) was used according to manufacturer’s instructions.

### qPCR

cDNA was quantified by real-time PCR using the SYBR green method with 2x SYBR Green PCR Master Mix (Thermo Fisher Scientific) and a StepOnePlus™ Real-Time PCR System (Thermo Fisher Scientific). qPCR primers are listed in Table 2. qPCR assays were run in duplicate. For each sample, the expression value for a gene of interest was first normalized for *Hprt* using the ddCt method and then divided by the mean of the control group to derive a relative gene expression value.

**Table 2:**
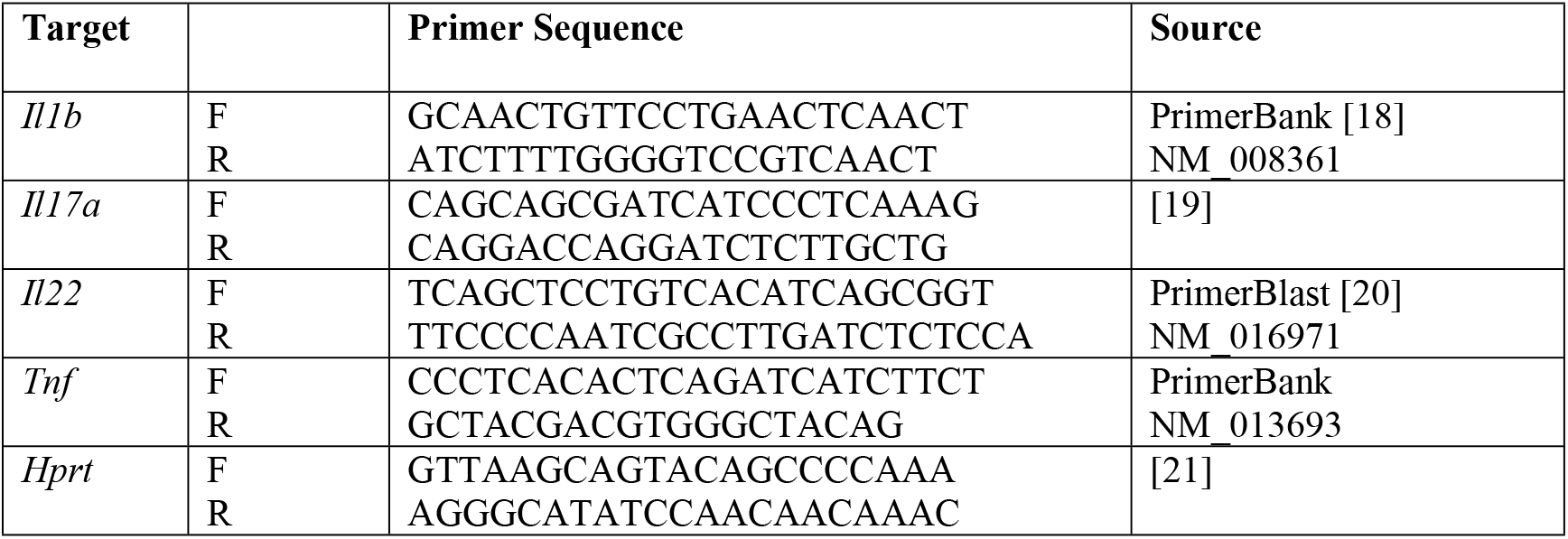
Primers for qPCR.

### ELISA

To measure serum IL-23 and IL-17A, Thermo Fisher Scientific ELISA kits was used according to manufacturer’s instructions.

### Micro-computed tomography (μCT)

Femurs were scanned using a μCT 35 (Scanco Medical AG). Scans were performed in 70% ethanol using a voxel size of 7 μm, X-ray tube potential of 55 kVp, intensity of 0.145 Ma, and integration time of 600 ms. A region beginning 0.35 mm proximal to the growth plate and extending 1 mm proximally was selected for trabecular bone analysis. A second region 0.6 mm in length and centered at the midpoint of the femur was used to calculate cortical bone parameters. The region of interest was thresholded using a global threshold that set the bone/marrow cutoff at 352.3 mg HA/cm^3^ for trabecular bone and 589.4 mg HA/cm^3^ for cortical bone. 3D microstructural properties of bone were calculated using software supplied by the manufacturer.

### Statistics

Graphing and statistical analysis were done with GraphPad Prism 8 software. The unpaired t-test was used to compare two experimental groups. Experiments with more than two groups were analyzed by one-way ANOVA followed by post hoc Tukey’s, with significance set at *p* < 0.05. Graphs show data points for individual mice and bars for mean ± standard deviation (SD) of the group.

## Results

### Hydrodynamic injection of IL-23 EEV in B10.RIII mice induces arthritis, colitis, and psoriasis-like skin disease

We hydrodynamically injected adult male B10.RIII mice with IL-23 EEV on day 0 and monitored disease development over 2 weeks. Control mice did not receive an injection. Clinical arthritis scores, paw swelling, and body weight were measured every other day. On day 14, the animals were euthanized, spleen and colon were weighed, and tissues were harvested for histology or further *in vitro* analysis. The general outline of this experiment is shown in Fig 1A and was used throughout this study.

**Fig 1.**
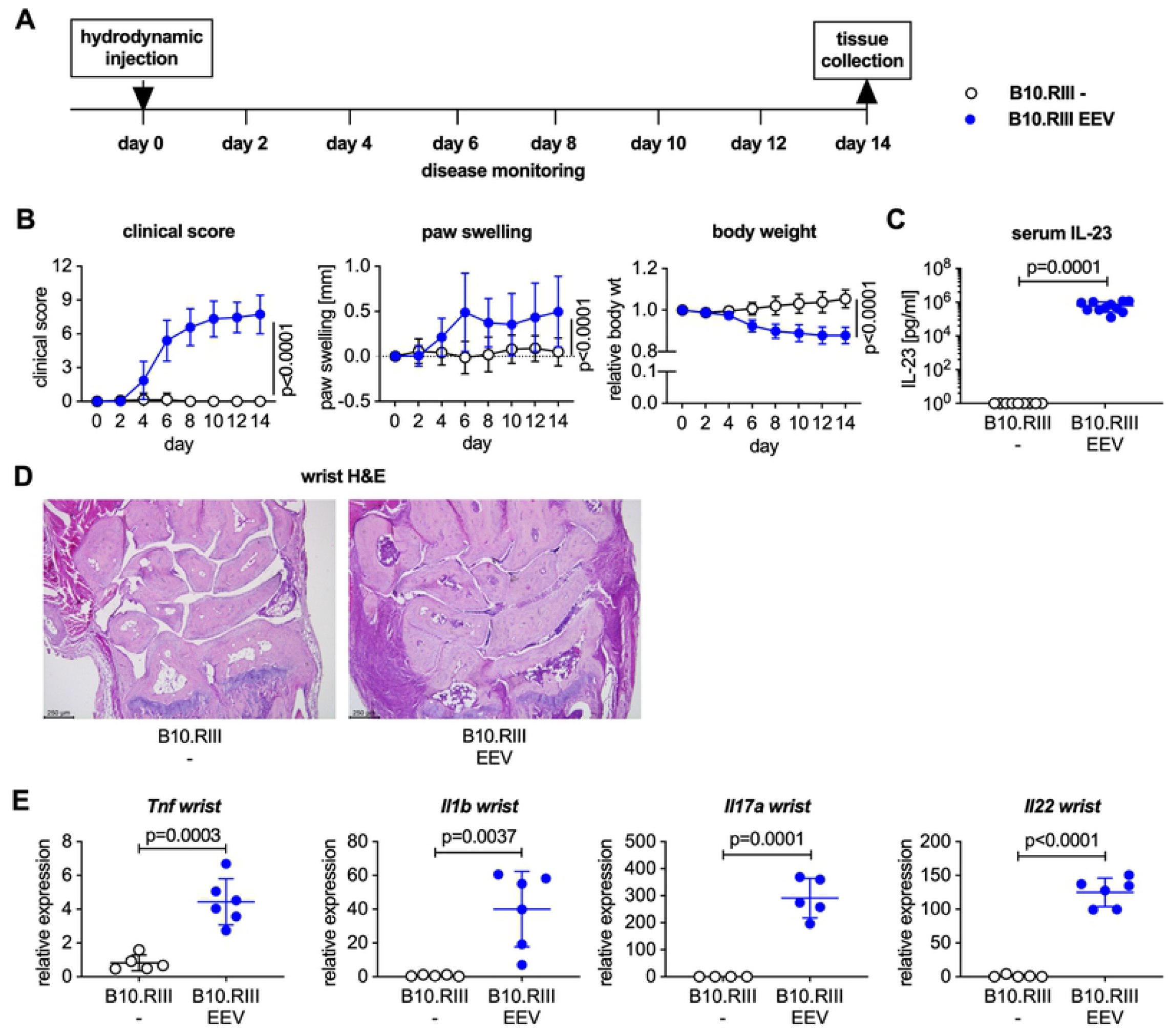
Male B10.RIII mice develop arthritis after hydrodynamic injection of IL-23 EEV. (A) Experimental layout for IL-23 EEV-induced arthritis experiments. (B) 8-12 week-old male B10.RIII mice received 500 ng IL-23 EEV on day 0 via hydrodynamic tail vein injection; control mice received no injection (n=12-15 per group). Clinical score (mean ± SD). Paw swelling (mean ± SD). Body weight (mean ± SD). (C) Serum IL-23 concentrations on day 14 were determined by ELISA (n=9-11 per group). (D) Representative H&E stained sections of day 14 wrists from control and IL-23 EEV injected mice, scale bar 250 μm. (E) Gene expression analysis in the wrists on day 14 (n=5-6 per group). qPCR data for individual samples were first normalized by *Hprt* expression and then divided by the mean of the uninjected control group. P values on day 14 were determined by unpaired t-test.

Upon injection of male B10.RIII mice with 500 ng IL-23 EEV, all animals developed clinical arthritis within 6 days (Fig 1B) with diffuse swelling of the forepaws, mid feet and ankles. IL-23 EEV injected mice also lost ~15% of their baseline body weight over 2 weeks (Fig 1B). This weight loss was the determining factor for choosing day 14 as study endpoint. Weight loss has not previously been reported in IL-23 EEV injected mice.

Serum IL-23 levels were significantly elevated in IL-23 EEV injected mice and below the level of detection in uninjected controls (Fig 1C). Representative H&E stained sections demonstrate a dense inflammatory infiltrate in the wrist of the IL-23 EEV injected mice compared to healthy controls (Fig 1D). Consistent with the observed clinical and histopathological arthritis phenotype, we found increased expression levels of *Tnf, Il1b* and the IL-23 target genes *Il17a* and *Il22* in the joints of IL-23 EEV injected mice (Fig 1E).

We also observed a statistically significant increase in colon weight compared to control mice (Fig 2A) suggesting the development of colitis. This was confirmed histologically. Compared to uninjected controls, H&E stained colon sections from IL-23 EEV injected mice demonstrated epithelial hyperplasia with crypt elongation, erosions of the epithelial surface and a mixed inflammatory infiltrate in the lamina propria (Fig 2B). Gene expression analysis by qPCR showed elevated expression of *Tnf, Il1b, Il17a*, and *Il22* in the colon of IL-23 EEV injected mice (Fig 2C). Consistent with previous reports [13–15, 22] we observed erythema and scaling of the skin of the ears and paws. H&E stained ear sections showed hyperplasia of the epithelial layer, thickening and inflammatory infiltration of the dermis (Fig 2D). Gene expression analysis in paw skin demonstrated increased expression of *Tnf, Il1b, Il17a*, and *Il22* in mice injected with IL-23 EEV (Fig 2E), consistent with the psoriasis-like changes seen on histology of the ears. Contrary to previous reports [13, 14], we did not find spinal inflammation on lumbar spine sections (Fig 2F) or any evidence for either posterior or anterior uveitis on ocular sections (Fig 2G) after IL-23 EEV injection. Together these data demonstrate that IL-23 overexpression in B10.RIII mice in our laboratory results in the development of arthritis, colitis, psoriasis-like skin disease but not spondylitis or uveitis.

**Fig 2.**
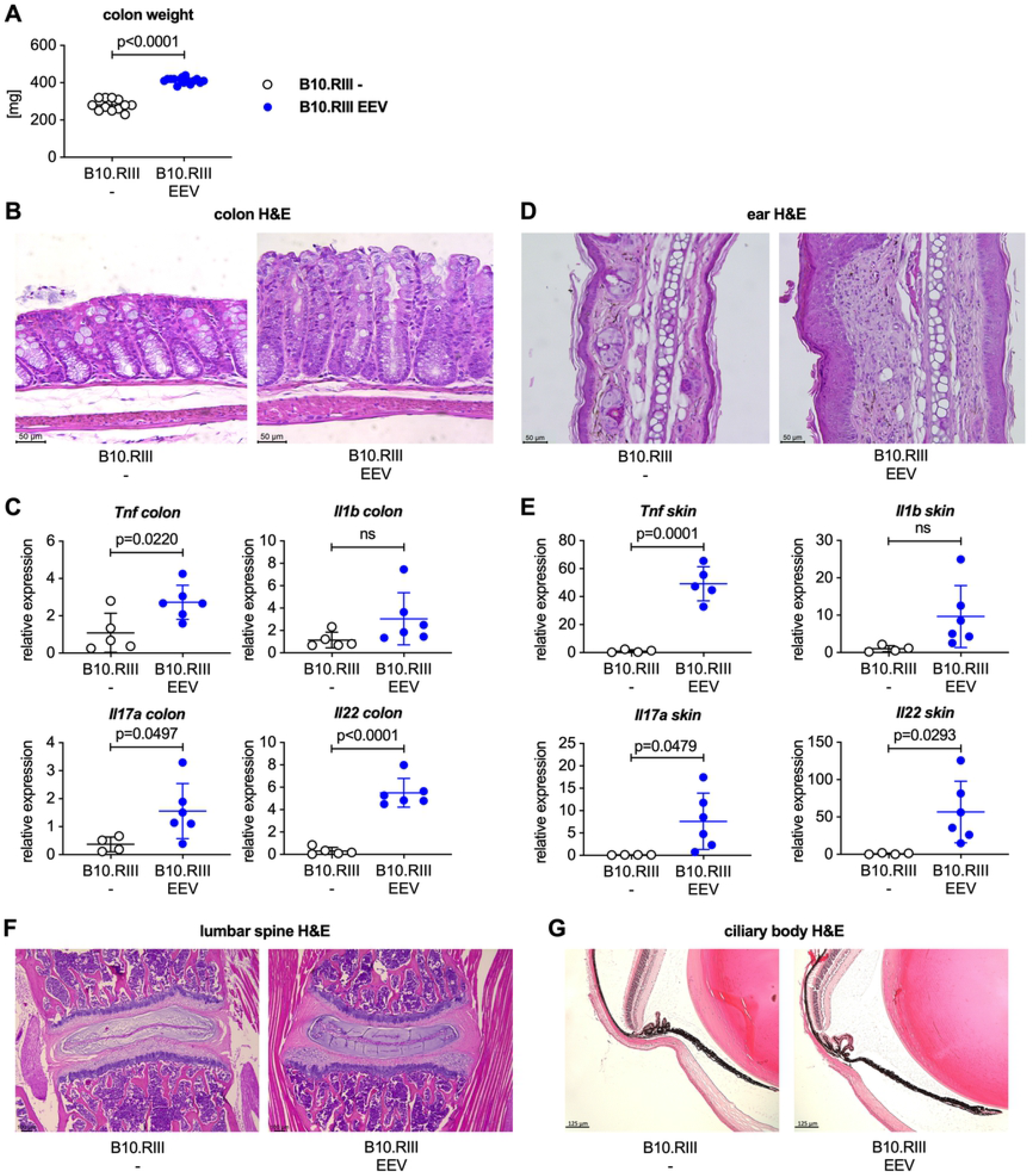
Male B10.RIII mice injected with IL-23 EEV develop colitis and psoriasis-like skin disease but not spondylitis or uveitis. (A) Colon weight of the animals in the experiment shown in Fig. 1. (B) Representative H&E stained colon sections from control and IL-23 EEV injected mice, scale bar 50 μm. (C) Expression of *Tnf, Il1b, Il17a*, and *Il22* in the colon was analyzed by qPCR (n=5-6 per group). (D) Representative H&E stained sections of day 14 ears from control and IL-23 EEV injected mice, scale bar 50 μm. (E) Expression for *Tnf, Il1b, Il17a*, and *Il22* in skin from the forepaws was analyzed by qPCR. P values were determined by unpaired t-test. (F) Representative H&E stained sections of day 14 lumbar spine from control and IL-23 EEV injected mice, scale bar 100 μm. (G) Representative H&E stained sections of day 14 eyes from control and IL-23 EEV injected mice, scale bar 125 μm.

IL-23 EEV injected mice had enlarged spleens compared with uninjected control animals (Fig 3A). Flow cytometric analysis of splenocytes (Fig 3B and Fig S1) demonstrated a significant influx of neutrophils, dendritic cells, and macrophages in the spleens of the IL-23 EEV injected mice, whereas the absolute numbers of B and T cells were not significantly different.

**Fig 3.**
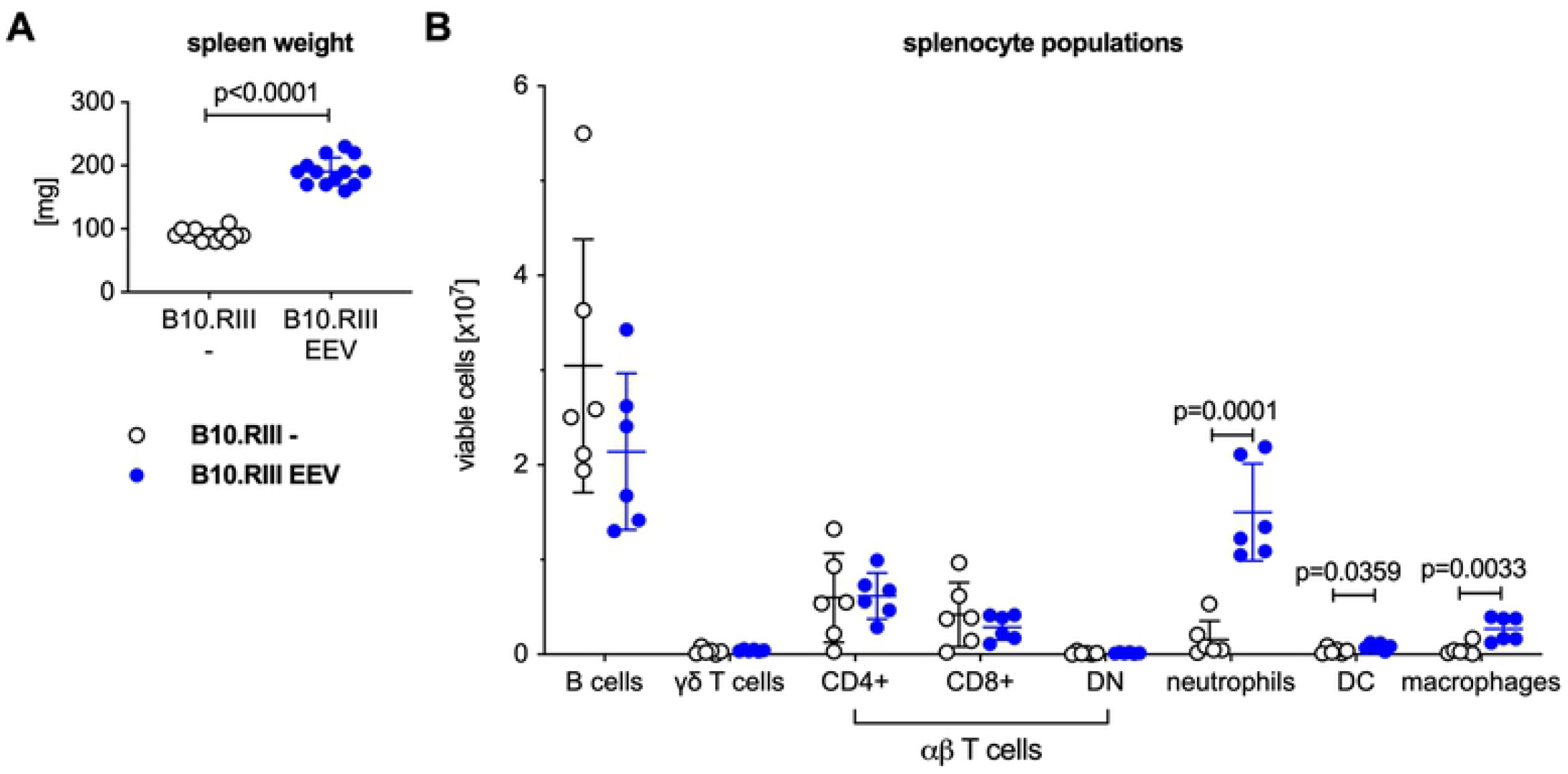
Systemic overexpression of IL-23 results in expansion of myeloid cells in the spleen. (A) Spleen weight of the mice from the experiment in Fig. 1. (B) For a subset of animals, single cell suspensions were prepared from the spleen and analyzed by flow cytometry, n=5 per group. P values were determined by unpaired t-test.

### Overexpression of IL-23 fails to induce arthritis in B6 mice, while extra-articular manifestations of IL-23-induced disease are present

In contrast to B10.RIII mice, male B6 mice injected with 500 ng IL-23 EEV did not develop clinical arthritis (Fig 4A). In fact, paw thickness of IL-23 EEV injected B6 mice decreased slightly over the 2-week period compared to controls. This was likely the result of weight loss (~15% over 2 weeks) and associated depletion of body fat in mice injected with IL-23 EEV, which was equally observed in B10.RIII and B6 mice (Fig 4A). B10.RIII and B6 mice had comparable serum levels of IL-23 and IL-17A on day 14 (Fig 4B). H&E stained sections of wrists confirmed the absence of arthritis in IL-23 EEV injected B6 mice compared to the susceptible B10.RIII strain (Fig 4C), and there was no significant induction of *Tnf, Il1b, Il17a* or *Il22* in the wrists of IL-23 EEV injected B6 mice (Fig 4D).

**Fig 4.**
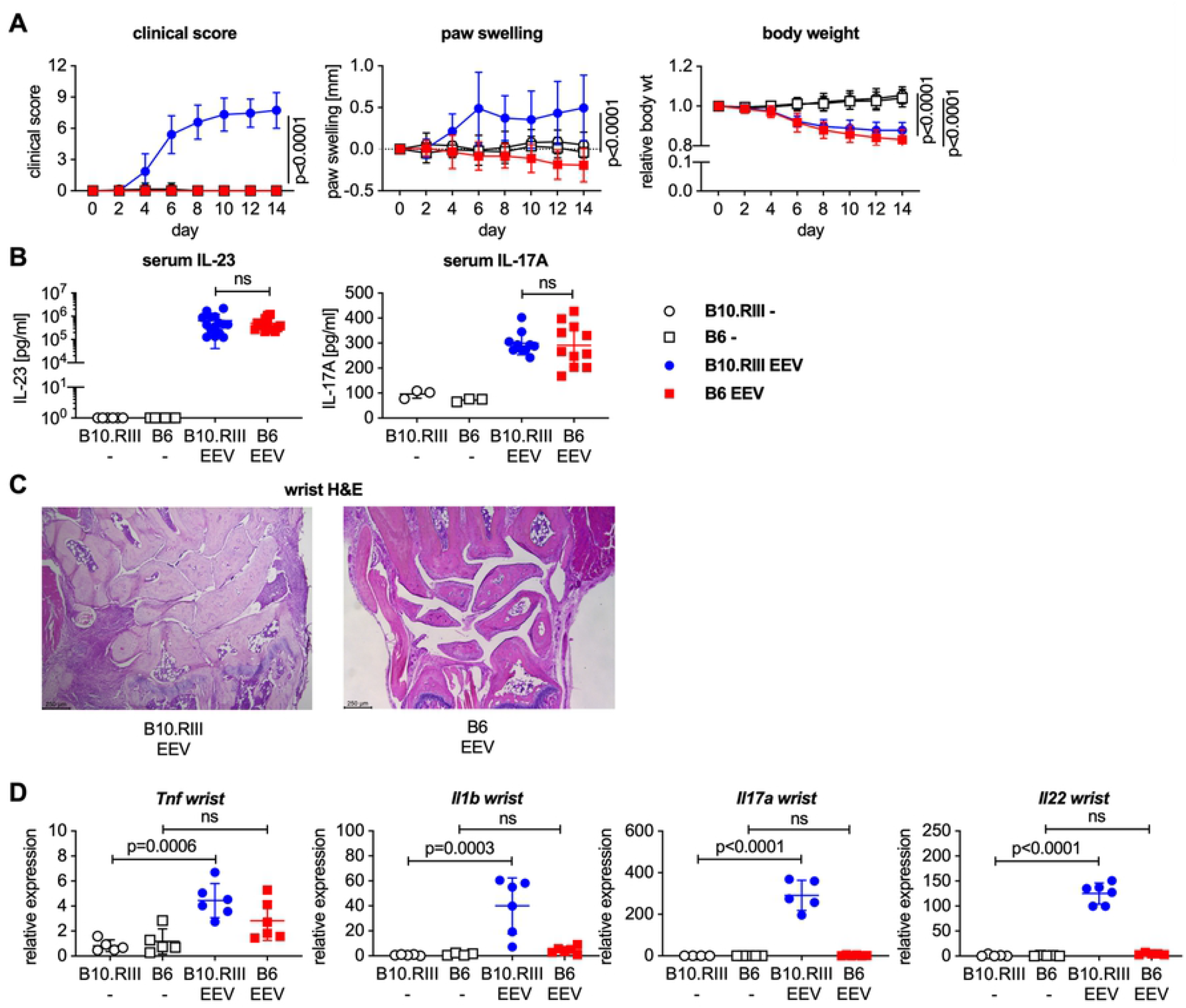
Male B6 do not develop arthritis after hydrodynamic injection of IL-23 EEV. 8-12 week-old male B10.RIII and C57BL/6 mice received 500 ng IL-23 EEV on day 0 via hydrodynamic tail vein injection; control mice received no injection (n=12-15 per group). (A) Clinical score (mean ± SD). Paw swelling (mean ± SD). Body weight (mean ± SD). (B) Serum IL-23 and IL-17A concentrations on day 14 were determined by ELISA. (C) Representative H&E stained sections of day 14 wrists from IL-23 EEV injected B10.RIII and B6 mice, scale bar 250 μm. (D) Gene expression analysis in the wrists on day 14 (n=5-6 per group). qPCR data for individual samples were first normalized by *Hprt* expression and then divided by the mean of all (B10.RIII and B6) uninjected control mice. P values were determined by unpaired t-test.

IL-23 EEV injected B6 mice developed colitis (Fig 5A-C) and psoriasis-like skin disease (Fig 5D and 5E), although these changes was less severe than in B10.RIII mice. Spleen enlargement after IL-23 EEV injection was also present but less severe in B6 mice compared with B10.RIII EEV mice (Fig 6A).

**Fig 5.**
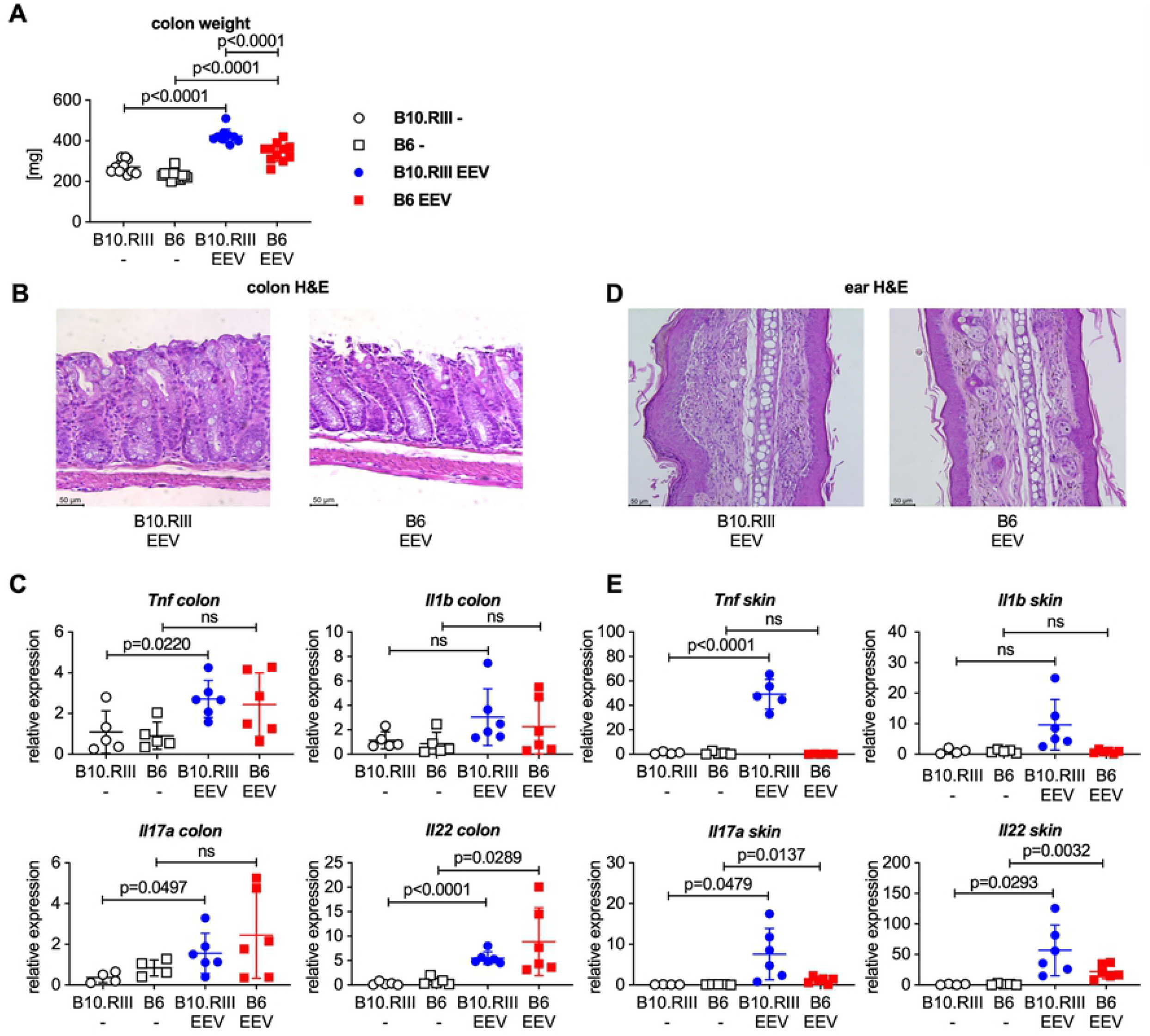
Male B6 mice develop colitis and mild psoriasis-like skin disease after hydrodynamic injection of IL-23 EEV. Data are from the animals in the experiment shown in Fig. 4. (A) Colon weight on day 14. (B) Representative H&E stained sections of day 14 colon from IL-23 EEV injected B10.RIII and B6 mice, scale bar 50 μm. (C) Gene expression for *Tnf, Il1b, Il17a*, and *Il22* in the colon was analyzed by qPCR. (D) Representative H&E stained sections of day 14 ears from IL-23 EEV injected B10.RIII and B6 mice, scale bar 50 μm. (E) Gene expression for *Tnf, Il1b, Il17a*, and *Il22* in forepaw skin was analyzed by qPCR.

**Fig 6.**
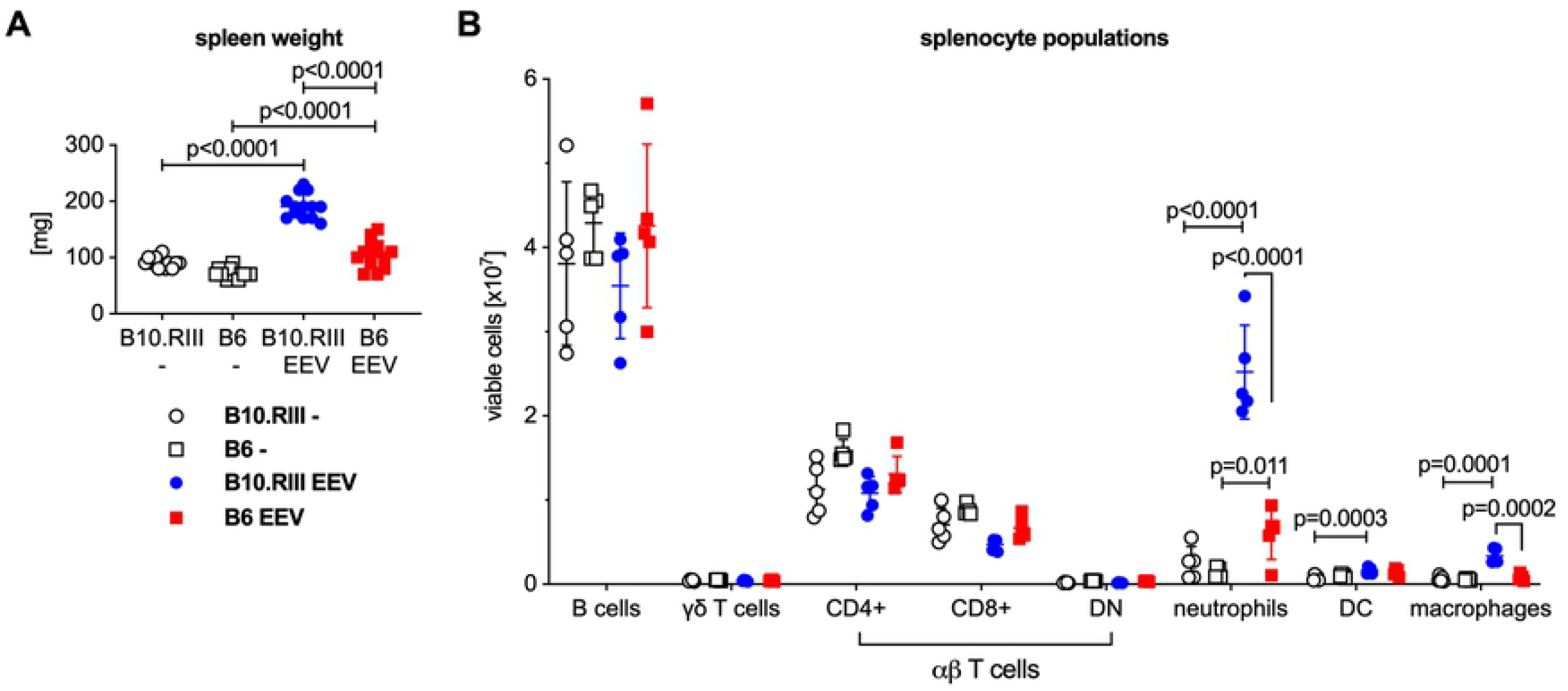
Male B6 mice accumulate significantly fewer myeloid cells than B10.RIII mice in the spleen after hydrodynamic injection of IL-23 EEV. (A) Spleen weight of the mice from the experiment in Fig. 4. (B) For a subset of animals, single cell suspensions were prepared from the spleens and analyzed by flow cytometry, n=5 animals per group. P values were determined by unpaired t-test.

Comparative flow cytometric analysis (Fig 6B) showed that the influx of myeloid cells after IL-23 EEV injection was less pronounced in B6 mice whereas lymphoid cell populations did not differ significantly.

The femurs of B10.RIII and B6 mice were analyzed by μCT two weeks after IL-23 EEV injection for inflammation-induced bone loss. Both strains had significantly less trabecular and cortical bone in their femurs after IL-23 EEV injection compared with uninjected littermates (Fig 7A). Representative images are shown in Fig 7B and 7C. Total mineral density did not change after IL-23 EEV injection. Additional μCT parameters are available for reference in Fig S2. Taken together these data demonstrate strain dependence of some IL-23 EEV induced phenotypes but not others. While male B6 mice did not develop arthritis despite similar serum IL-23 and IL-17A levels, other disease manifestations (weight loss, colitis, psoriasis-like skin disease, bone loss) were observed in both strains.

**Fig 7.**
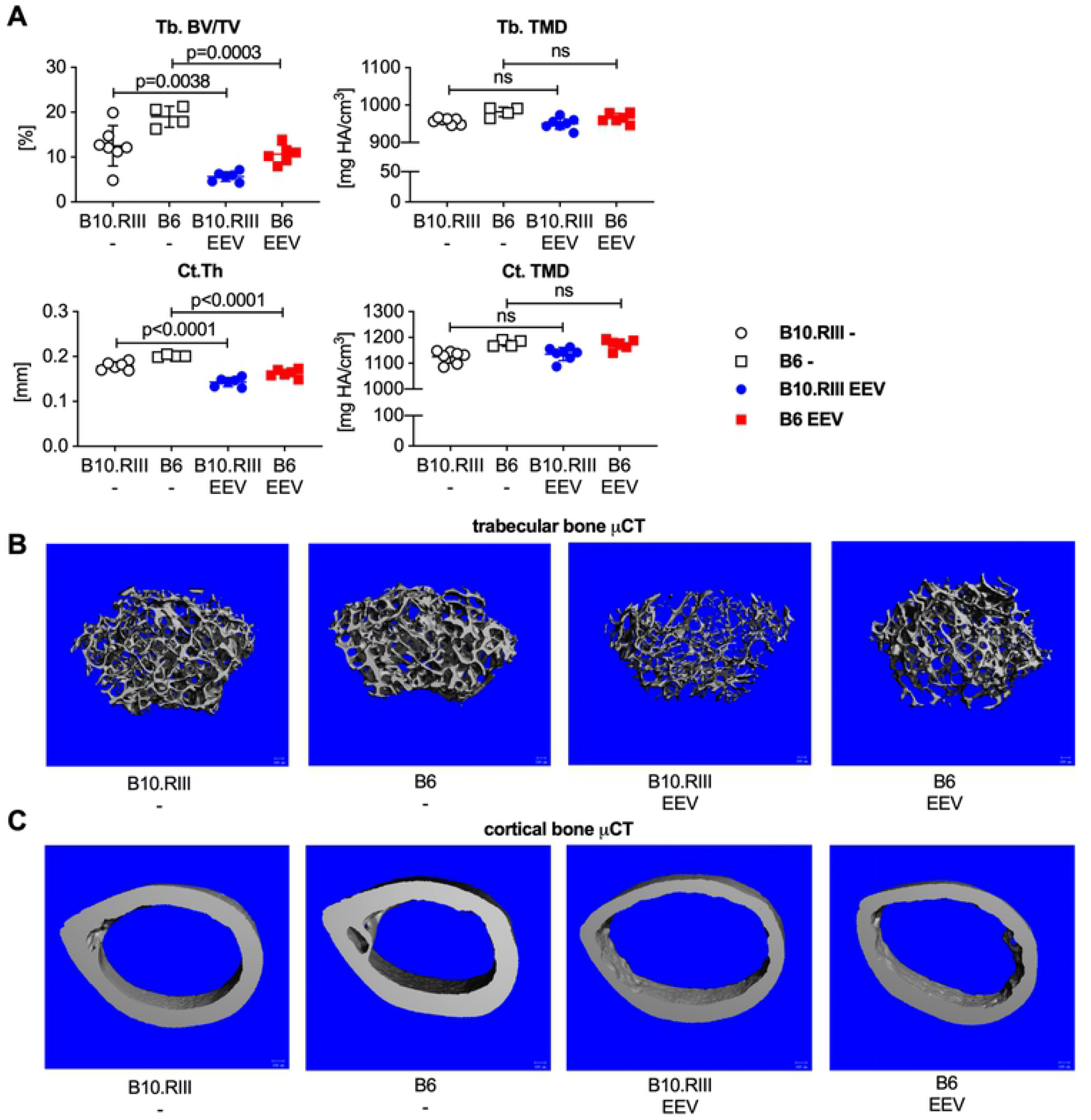
Both B10.RIII and B6 mice lose trabecular and cortical bone after IL-23 EEV injection. (A) μCT trabecular and cortical bone parameters from femurs of control and IL-23 EEV injected B10.RIII and B6 14 days after IL-23 EEV injection (n=4-7 per group). Trabecular bone volume/total volume (Tb. BV/TV), trabecular bone total mineral density (Tb. TMD), cortical thickness (Ct. Th), and cortical total mineral density (Ct. TMD). P values were determined by unpaired t-test. (B) Representative μCT images of trabecular bone and (C) representative μCT images of cortical bone from control and IL-23 EEV injected B10.RIII and B6 mice.

### Overexpression of IL-23 results in more severe arthritis in female B10.RIII mice compared to male B10.RIII mice

Male and female B10.RIII mice injected with IL-23 EEV developed clinical arthritis with 100% penetrance regardless of sex (Fig 8A). Clinical arthritis scores and paw swelling were significantly higher in female than male B10.RIII mice injected with IL-23 EEV (Fig 8A). Interestingly, female B10.RIII mice injected with IL-23 EEV did not lose weight over the course of the two-week experiment (Fig 8A), and serum IL-23 levels were significantly lower in females than males despite receiving the same dose of IL-23 EEV (Fig 8B).

**Fig 8.**
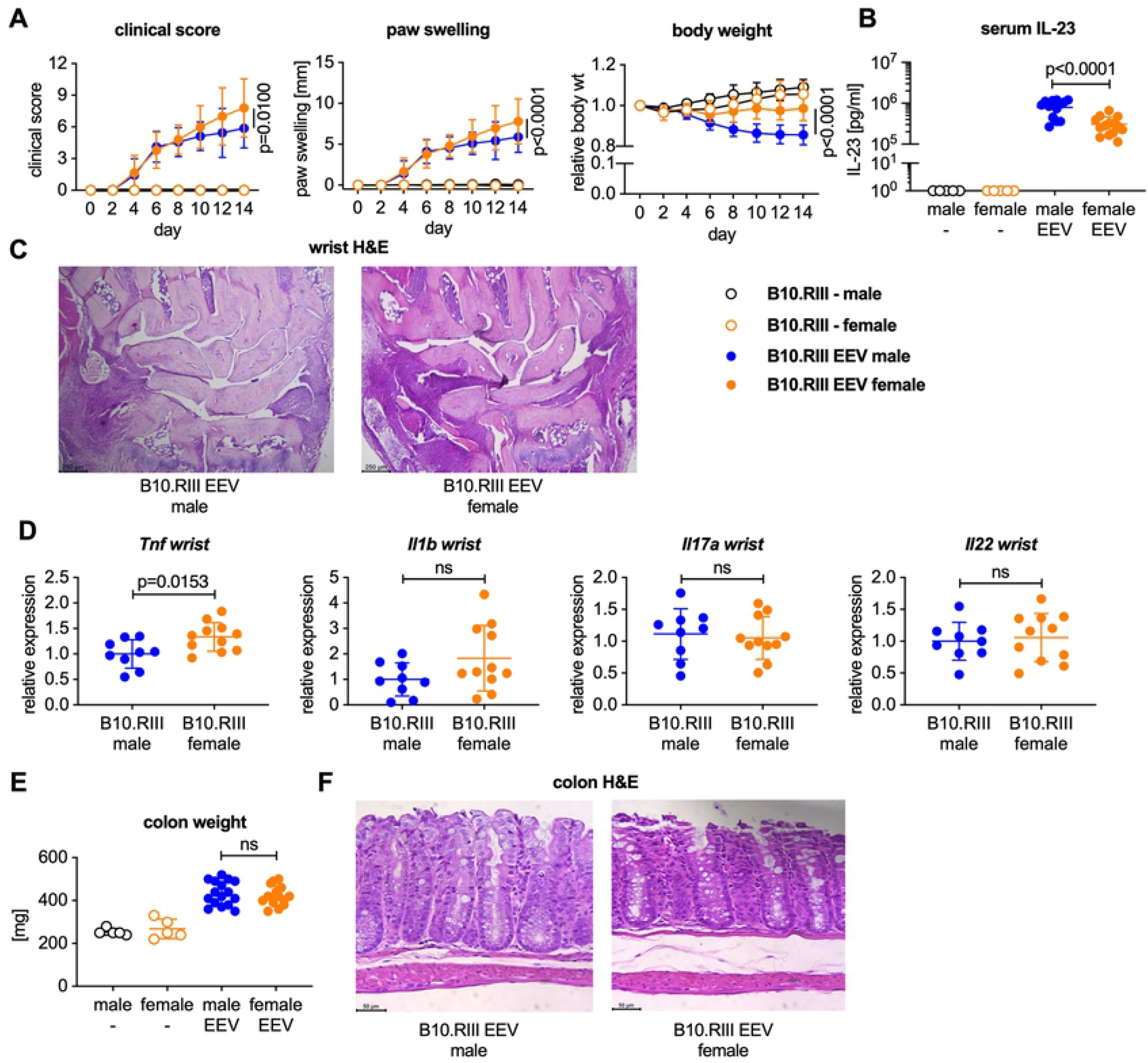
Female B10.RIII mice develop more severe arthritis than males but do not lose weight after hydrodynamic injection of IL-23 EEV. 8-12 week-old male and female B10.RIII mice received 500 ng IL-23 EEV on day 0 via hydrodynamic tail vein injection; control mice received no injection (n=14 in IL-23 EEV injected groups, n=5 in control groups). (A) Clinical score (mean ± SD). Paw swelling (mean ± SD). Body weight (mean ± SD). (B) Serum IL-23 on day 14 was determined by ELISA. (C) Representative H&E stained sections of day 14 wrists, scale bar 250 μm. (D) Gene expression analysis in the wrists on day 14 (n=9-11 per group). qPCR data for individual samples were first normalized by *Hprt* expression and then divided by the mean of the IL-23 EEV injected males. P values were determined by unpaired t-test. (E) Colon weight on day 14. (F) Representative H&E stained colon sections from male and female B10.RIII IL-23 EEV injected mice, scale bar 50 μm.

H&E stained sections showed inflammatory cells infiltration in the wrists of both male and female mice on day 14 after IL-23 EEV injection with higher levels of *Tnf* but not *Il1b, IL17a* or *Il22* in the female wrists compared to male wrists (Fig 8C and 8D). Female mice displayed a comparable increase in colon weight suggesting a similar degree of colitis relative to male mice after IL-23 EEV injection (Fig 8E and 8F).

Female mice had significantly larger spleens associated with a more pronounced influx of myeloid cells, compared to male mice on day 14 after IL-23 EEV injection (Fig 9A and 9B). Collectively, these data show that IL-23 overexpression results in distinct phenotypic differences between male and female B10.RIII mice; female mice had more severe arthritis and did not lose weight despite similar colitis severity.

**Fig 9.**
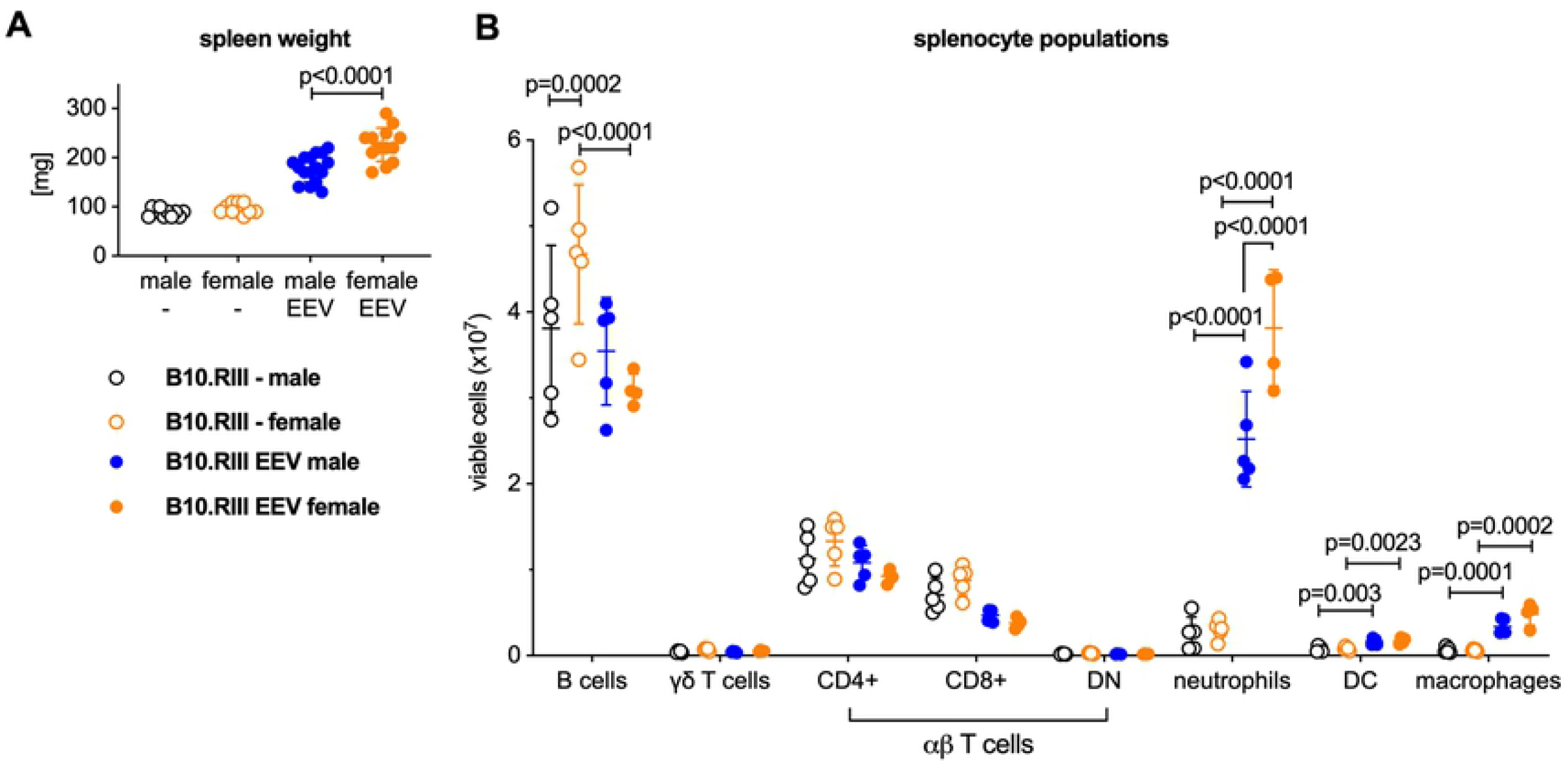
Female B10.RIII mice exhibit greater expansion of myeloid cells in the spleen after hydrodynamic injection of IL-23 EEV. (A) Spleen weight of the mice from the experiment in Fig.8. (B) For a subset of animals, single cell suspensions were prepared from the spleens and analyzed by flow cytometry, n=4-5 animals per group. P values were determined by unpaired t-test.

### Overexpression of IL-23 by injection 50 ng IL-23 EEV results in similar clinical scores and less weight loss than mice injected with 500 ng

Lastly, we hydrodynamically injected adult male B10.RIII mice with 5, 50, or 500 ng of IL-23 EEV to determine the impact of EEV dose on disease severity. Arthritis development after injection of 50 and 500 ng of IL-23 EEV was comparable but recipients of the 50 ng dose lost significantly less body weight over 14 days (Fig S3A). All 3 doses resulted in significant induction of serum IL-23 with a linear relationship between minicircle dose and IL-23 serum level (Fig S3 B). However, mice injected with 5 ng did not develop arthritis, weight loss, colitis or spleen enlargement (Fig S3 A-D) suggesting the existence of a threshold level of serum IL-23 that is required for the development of IL-23 EEV induced disease.These results indicate that an IL-23 minicircle dose of 50 ng in male B10.RIII mice is optimal resulting in arthritis development without the severe weight loss observed with a 500 ng dose. This will permit studying the musculoskeletal phenotypes in IL-23 EEV injected mice in longer-term experiments.

## Discussion

We report here that IL-23-induced disease in B10.RIII mice in our laboratory is characterized by peripheral arthritis, psoriasis-like skin disease, colitis, weight loss, osteopenia and absence of spondylitis and uveitis. Male B6 mice injected with IL-23 EEV did not develop arthritis despite comparable serum IL-23 and IL-17A levels, degree of weight loss, and osteopenia with B10.RIII mice. Some of the IL-23-induced phenotypes in B10.RIII mice exhibited sexual dimorphism in that female B10.RIII mice had more severe arthritis than males but did not lose weight.

The rapid disease onset of arthritis and development of psoriasis-like skin changes in B10.RIII mice injected with IL-23 EEV is consistent with prior studies [13–15, 23]. However, we did not see spondylitis or uveitis which had been observed by others [13, 14]. On the other hand, B10.RIII mice in our lab developed colitis and lost weight with higher doses of IL-23 EEV, which had not been seen previously. Two studies investigated gastrointestinal phenotypes in IL-23 minicircle injected B6 mice: Paustian et al. reported IL-23 induced small bowel inflammation whereas Chan et al. stated the absence of inflammation in thew small intestine [24, 25]. The reasons for these discrepancies between studies are not clear. One potential explanation involves differences in the intestinal microbiota between animal facilities [26].It is also possible that difference in the IL-23 expression vector contribute to the observed phenotypic differences. We used the IL-23 enhanced episomal vector from System Biosciences. Others have used IL-23 minicircles from the same vendor or prepared IL-23 minicircles themselves. The EEV vector is extremely efficient, leading to arthritis induction with as little as 50 ng of EEV plasmid. Differences in promoter sequence and kinetics of IL-23 induction may play a role in determining the disease phenotype.We found that weight loss was correlated with serum IL-23 levels (and EEV dose) and could reflect effects on IL-23 receptor expressing lymphocytes in adipose tissue that only become apparent with higher levels of serum IL-23 [27].

Strain dependence of arthritis phenotypes has been reported for many mouse arthritis models. B10.RIII mice are highly susceptible to collagen induced arthritis (CIA). The B10.RIII strain is congenic for the RIII-derived H-2^r^ region, a variant of the mouse MHC that is permissive for productive immunization with a variety of type 2 collagens [28]. However, B10.RIII mice are also highly susceptible to collagen antibody-induced arthritis (CAIA) which bypasses the adaptive immune response required for the formation of pathogenic collagen-specific antibodies and reads out “down-stream” effector mechanisms [29, 30] while the RIII strain, which has the same H-2^r^ haplotype as B10.RIII is not susceptible to CAIA [29]. This suggests that the region(s) determining differential arthritis susceptibility between B10.RIII (H-2^r^) and B6 (H-2^b^) mice in the IL-23 model could be anywhere in the genome.

One potential lead to understanding the differential susceptibility to IL-23 induced arthritis could be the observed splenomegaly driven by an expansion of neutrophils, dendritic cells, and macrophages in mice injected with IL-23 EEV. There was a correlation between the severity of splenomegaly and arthritis development both in the comparison of male B10.RIII and B6 mice and of male and female B10.RIII mice injected with IL-23 EEV. This is consistent with previous reports describing that in IL-23–induced arthritis, the expansion and activity of myeloid cells was crucial for arthritis development. Adamopolous et al. depleted myeloid cells with clodronate, significantly decreasing disease severity and paw swelling [23]. The same group identified that spleen neutrophil counts were increased in mice injected with IL-23 minicircles. When treated with anti-γδTCR antibodies, there was no significant neutrophil expansion in the spleen which correlated with reduced arthritis severity [15].

The NIH considers sex as a biological variable to be critical for validation and generalizability of research findings [17]. We observed that overexpression of IL-23 resulted in more severe arthritis in female B10.RIII mice compared to male B10.RIII mice. Increased susceptibility to inflammation does not appear to be an inherent feature of female B10.RIII mice as a study in OVA-induced allergic airway inflammation found no difference between male and female B10.RIII mice [31]. A potential area for investigation is that females have significantly more adipose tissue than males, and adipose tissue macrophages have been found to be a potent source of inflammatory cytokines {Giles, 2018 #11439}.

A limitation of this study is that the mechanisms regulating the strain and sex-specific differences in arthritis susceptibility remain unclear. The absence of arthritis in B6 mice injected with IL-23 EEV reduces the utility of this model for mechanistic studies as most genetically modified mice have been generated on a B6 genetic background. However, a strength of our study is the delineation of highly robust and reproducible phenotypes which provides opportunities for further explorations of SpA pathogenesis including the impact of genetic polymorphisms, intestinal microbiota and sex. Moreover, compared with CAIA or the K/BxN serum transfer model the IL-23 EEV model is relatively cheap.

In summary, hydrodynamic injection of IL-23 EEV in B10.RIII mice is a robust, time-efficient and economical model for the study of SpA pathogenesis. The differential susceptibility of B6 mice with regard to the development of arthritis and extra-articular manifestations suggests organ-specific genetic control mechanisms of IL-23 driven inflammation. The mechanisms behind these differences and the sexual dimorphism observed in this study may be relevant to the understanding of human SpA and warrant further investigation.

## Acknowledgements

Support for this research was provided by grants from NIAMS (R03AR066357-01A1), the Rheumatology Research Foundation, and the Evergreen Fund. Histology services were provided by the Center for Skeletal Research (CSR), an NIH funded program (P30 AR075042) supported by NIAMS. The Research to Prevent Blindness supports the Casey Eye Institute at Oregon Health & Science University.

## Supporting information

**S1 Fig.**
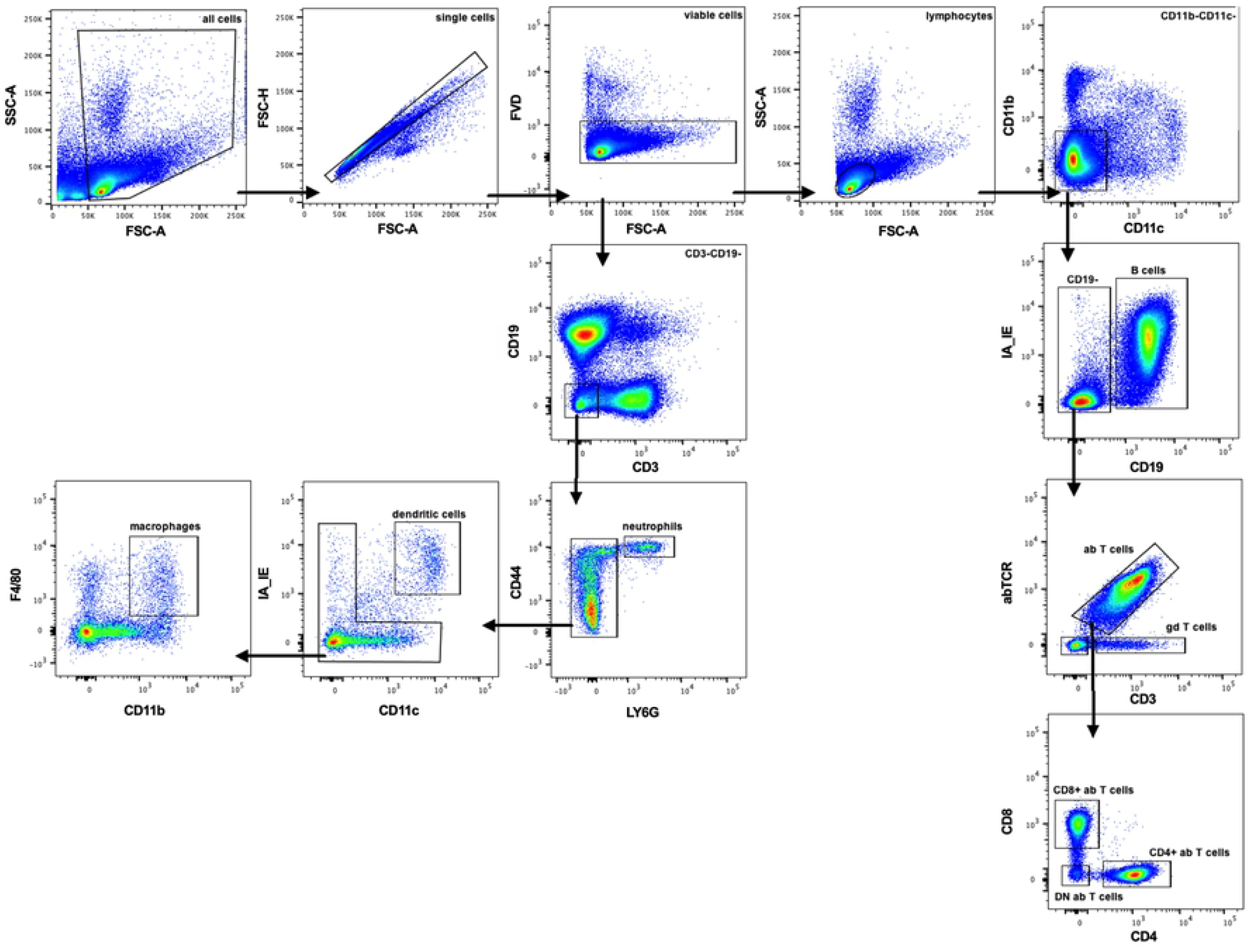
Gating strategy for flow-cytometric analysis of single cell suspensions. Cell analysis was performed on BD LSRFortessa. Gating was performed using FlowJo software.

**S2 Fig.**
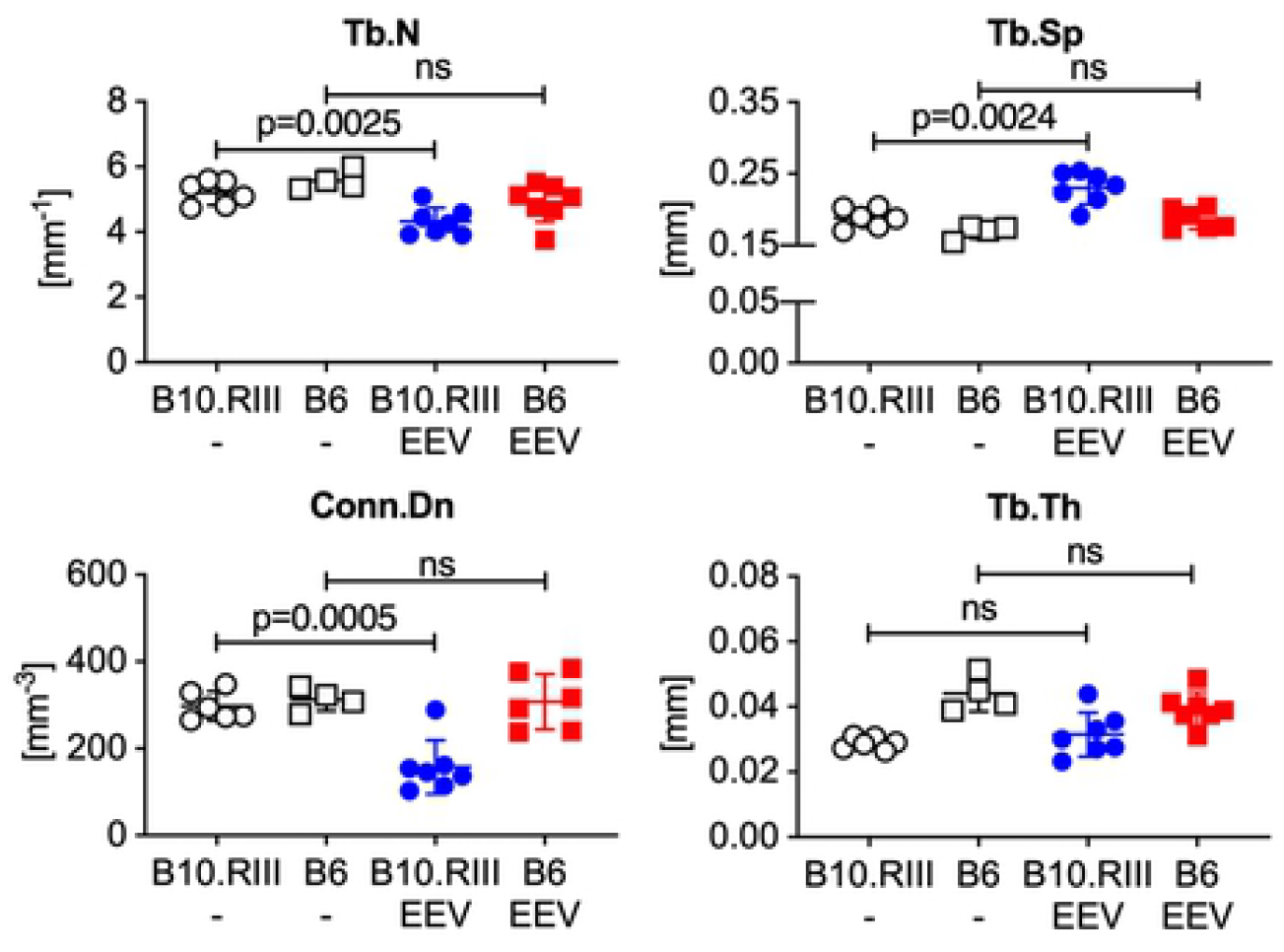
Additional μCT bone parameters. μCT trabecular bone parameters from femurs of control and IL-23 EEV injected B10.RIII and B6 14 days after IL-23 EEV injection (n=4-7 per group). Trabecular number (Tb. N), trabecular spacing (Tb. Sp), connective density (Conn.Dn), and trabecular thickness (Tb.Th). P values were determined by unpaired t-test.

**S3 Fig.**
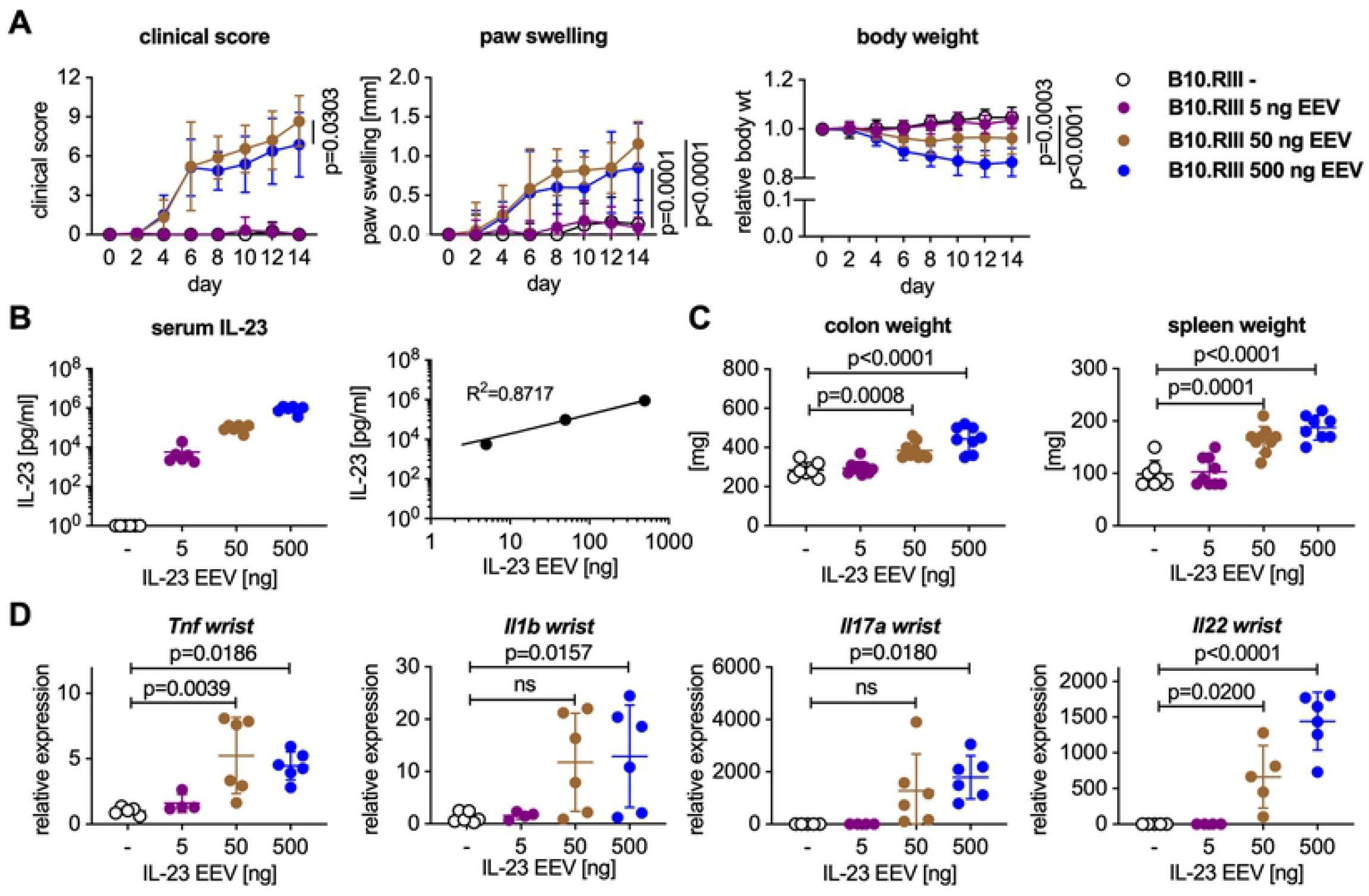
IL-23 EEV *in vivo* dose titration. 8-12 week-old male mice received 5, 50, or 500 ng IL-23 EEV on day 0 via hydrodynamic tail vein injection; control mice received no injection (n=7-8 animals per group). (A) Clinical score (mean ± SD), paw swelling (mean ± SD), body weight (mean ± SD). (B) Serum IL-23 on day 14 was determined by ELISA. (C) Spleen weight, colon weight. Dotted lines represent the mean weights for control mice. (D) Gene expression analysis in the wrists on day 14. qPCR data for individual samples were first normalized by *Hprt* expression and then divided by the mean of the uninjected control group.

